# On the evolutionary ecology of multidrug resistance in bacteria

**DOI:** 10.1101/233957

**Authors:** Sonja Lehtinen, François Blanquart, Marc Lipsitch, Christophe Fraser, The Maela Pneumococcal Collaboration

**Affiliations:** The Oxford Big Data Institute, Nuffield Department of Medicine, University of Oxford, Oxford, UK; Center for Interdisciplinary Research in Biology, College de France, Paris, France; Department of Infectious Disease Epidemiology, Imperial College London, London, UK; Center for Communicable Disease Dynamics, Harvard T. H. Chan School of Public Health, Boston, USA; Departments of Epidemiology and Immunology and Infectious Diseases, Harvard T. H. Chan School of Public Health, Boston, USA; Stephen Bentley (The Wellcome Trust Sanger Institute, Cambridge, UK), Nicholas J. Croucher (Imperial College London, London, UK), John Lees (Wellcome Trust Sanger Institute, Cambridge, UK; NYU School of Medicine, New York, USA), Paul Turner (Centre for Tropical Medicine and Global Health, Nuffield Department of Medicine, University of Oxford, UK)

## Abstract

Resistance against different antibiotics appears on the same bacterial strains more often than expected by chance, leading to high frequencies of multidrug resistance. There are multiple explanations for this observation, but these tend to be specific to subsets of antibiotics and/or bacterial species, whereas the trend is pervasive. Here, we consider the question in terms of strain ecology: explaining why resistance to different antibiotics is often seen on the same strain requires an understanding of the competition between strains with different resistance profiles. This work builds on models originally proposed to explain another aspect of strain competition: the stable coexistence of antibiotic sensitivity and resistance observed in a number of bacterial species. We first demonstrate a partial structural similarity in these models of coexistence. We then generalise this unified underlying model to multidrug resistance and show that models with this structure predict high levels of association between resistance to different drugs and high multidrug resistance frequencies. We test predictions from this model in six bacterial datasets and find them to be qualitatively consistent with observed trends. The higher than expected frequencies of multidrug resistance are often interpreted as evidence that these strains are out-competing strains with lower resistance multiplicity. Our work provides an alternative explanation that is compatible with long-term stability in resistance frequencies.

**Author summary:** Antibiotic resistance is a serious public health concern, yet the ecology and evolution of drug resistance are not fully understood. This impacts our ability to design effective interventions to combat resistance. From a public health point of view, multidrug resistance is particularly problematic because resistance to different antibiotics is often seen on the same bacterial strains, which leads to high frequencies of multidrug resistance and limits treatment options. This work seeks to explain this trend in terms of strain ecology and the competition between strains with different resistance profiles. Building on recent work exploring why resistant bacteria are not out-competing sensitive bacteria, we show that models originally proposed to explain this observation also predict high multidrug resistance frequencies. These models are therefore a unifying explanation for two pervasive trends in resistance dynamics. In terms of public health, the implication of our results is that new resistances are likeliest to be found on already multidrug resistant strains and that changing patterns of prescription may not be enough to combat multidrug resistance.

## 1 Introduction

Antibiotic resistance and, in particular, multidrug resistance (MDR) are public health threats. Multidrug resistant infections are associated with poorer clinical outcomes and higher cost of treatment than other infections [1, 2] and there is concern that the emergence of pan-resistant strains (pathogens resistant to all available antibiotics) will render some infections untreatable [3].

From the point of view of finding effective treatment options, multidrug resistance is particularly problematic because resistance to different antibiotics tends to be concentrated on the same strains: positive correlations between resistance to different drugs have been found in multiple species (including *Streptococcus pneumoniae, Neisseria gonorrhoeae, Staphylococcus aureus, Escherichia coli, Klebsiella pneumoniae, Pseudomonas aeruginosa* and *Mycobacterium tuberculosis*) [2]. In other words, the frequency of MDR strains is higher than we would expect from the frequencies of individual resistance determinants if these were distributed randomly in the population (‘MDR over-representation’).

Understanding the causes of this MDR over-representation is important for limiting the impact of resistance. A number of possible explanations have been suggested (Table 1) [2], but the extent to which these processes contribute to the trend remains uncertain. Many of the proposed mechanisms are specific to subsets of antibiotics and/or species. The pattern of MDR over-representation, on the other hand, is pervasive: correlations have been observed between resistance to antibiotics acting through different mechanisms, and between chromosomal and mobile genetic element (MGE) associated resistance determinants [2]. Explanations for MDR over-representation must therefore be either sufficiently general or sufficiently diverse to account for this pervasiveness.

**Table 1:** Processes which may contribute to MDR over-representation..

| Process | Notes |
| --- | --- |
| Shared resistance mechanisms | Particularly relevant for antibiotics of the same class (e.g. β-lactamases and some penicillin-binding protein mutations conferring resistance against multiple β-lactams), but also applicable for some drugs of different classes: there are examples of efflux pumps acting on multiple drugs in numerous species [4] and evidence for clinical relevance of efflux pumps in multidrug resistance [5]. However, MDR over-representation is also observed where efflux pumps are not thought to be a major mechanism of resistance (e.g. between β-lactams and other classes of antibiotics in *S. pneumoniae* [2]). |
| Linkage between resistance genes (when resistance is associated with particular alleles) | In discussing linkage between resistance genes (i.e. resistance genes being inherited together), it is helpful to distinguish between resistance mechanisms where a particular allele of a gene confers resistance (e.g. changes to the protein targeted by the antibiotics) and those where resistance is associated with the presence of a resistance gene (e.g. enzymes that break down the drug). When resistance is associated with specific alleles, genetic linkage is not a plausible mechanism for generating MDR over-representation: *a priori*, linkage is no more likely to promote association between resistance determinants than association between resistance to one antibiotic and sensitivity to another. |
| Linkage between resistance genes (when resistance is associated with presence of gene) | When resistance is associated with presence of a particular gene, absence of the gene from a particular MGE does not necessarily imply sensitivity to the antibiotic (the gene may be present on another element). As a consequence, spread of the MGE will spread resistance for resistance genes present on the element, but not spread sensitivity when resistance genes are absent. Linkage would thus favour association between resistance determinants. Even in this case, however, we would expect the effect of linkage to be transient: recombination and mutation eventually eliminate resistance determinants which do not confer a fitness advantage, even from mobile genetic elements. For example, in the PMEN1 pneumococcal lineage, there is evidence for loss of aminoglycoside resistance from the Tn916 transposon which encodes tetracycline, and sometimes macrolide, resistance [6]. |
| Correlated drug exposure of individual host | Correlated drug exposure could arise either through use of combination therapy or through sequential drug exposure if the first choice of drug fails. In practice, however, it is unclear how significant a role these mechanisms play in MDR over-representation. In the United Kingdom for example, monotherapy accounts for 98% of primary care prescriptions [7]. It is unclear whether treatment failure rates (below 20% for the most commonly prescribed antibiotics in UK primary care [7]) and patterns of subsequent drug prescription are enough to drive selection for multidrug resistance. |
| Cost epistasis (lower than expected fitness cost when multiple resistance determinants are present) | There is evidence of cost epistasis between resistance determinants occurring in laboratory competition experiments for some antibiotics (e.g. between streptomycin and rifampicin resistance in *Pseudomonas aeruginosa* [8] and in *E. coli* [9]; streptomycin and quinolone resistance in *E. coli* [10]; and rifampicin and ofloxacin resistance in *Mycobacterium smegmatis* [11]). Furthermore, for plasmid-associated resistance genes, cost epistasis could also arise if the presence of the plasmid in itself incurs a significant fitness cost (rather than the fitness cost depending on the specific resistance genes it carries). However, the extent to which epistasis plays a role *in vivo* remains unclear [12]. In particular, we would not, *a priori*, expect to observe cost epistasis between resistance determinants operating through entirely different mechanisms. |

In this paper, we approach the problem of explaining MDR over-representation in terms of strain ecology: explaining why resistance to different antibiotics is often seen on the same strain requires an understanding of the competition between strains with different resistance profiles. For models of such competition to be credible, they must capture observed trends in resistance dynamics whilst being ecologically plausible. Developing models that fulfil these criteria has not been trivial: sensitive and resistant strains compete for the same hosts and simple models of competition therefore predict that the fitter strain will out-compete the other (‘competitive exclusion’) [13]. However, this is rarely observed: resistance frequencies have remained intermediate over long time periods in a number of species. For example, sustained intermediate resistance frequencies are observed in Europe for various antibiotics and numerous species, including *E. coli, S. aureus* and *S. pneumoniae* (European Centre for Disease Prevention and Control Surveillance Atlas, available at https://atlas.ecdc.europa.eu). Stable coexistence is also observed in surveillance data from multiple other locations (Centre for Disease Dynamics, Economics and Policy, available at https://resistancemap.cddep.org/AntibioticResistance.php). For further review of evidence for stable coexistence, see references [13, 14].

Recent work has explored the role of i) host population structure [15, 16, 17], ii) pathogen strain structure [15, 14] and iii) within-host dynamics [18] in maintaining the coexistence of antibiotic sensitivity and resistance. In this paper, we identify a structural similarity in the first two categories of model: in these models, coexistence is maintained by mechanisms that introduce heterogeneity in the fitness benefit gained from resistance within the pathogen population. We show that models with this structure also predict high levels of association between resistance to different antibiotics: all resistance determinants will tend to be found where the fitness benefit gained from resistance is the greatest. The observed high frequency of multi-drug resistance is therefore in line with ecologically plausible models of coexistence, making these models a parsimonious explanation for both trends.

## 2 Results

### 2.1 Heterogeneity in the fitness effect of resistance: a generalised model of co-existence

In this section, we discuss competitive exclusion and previously proposed coexistence mechanisms in the context of multidrug resistance. We identify a structural similarity in plausible models of coexistence [16, 15, 14] and show that, in a multidrug context, models with this structure predict MDR over-representation. The model we present captures the dynamics of a bacterial species which is mostly carried asymptomatically (e.g. *E. coli, S. aureus* or *S. pneumoniae*), so the probability of a host being exposed to antibiotics does not depend on whether the host is infected with the pathogen [19]. Key results, however, are also applicable when this is not the case (see Discussion).

#### 2.1.1 Competitive exclusion in single and multi-drug systems

The coexistence of antibiotic sensitivity and resistance has been previously discussed in the context of competition between two strains (sensitive and resistance strains [13, 15, 14] or two resistant strains with different resistance profiles [16]). Simple models of such competition predict competitive exclusion [13]; we start by briefly re-introducing this result and then demonstrate that competitive exclusion also applies in a multidrug context.

We consider a SIS (susceptible-infectious-susceptible) model of resistant and sensitive variants of an otherwise genetically homogeneous pathogen (one strain) circulating in a homogeneous host population. To avoid ambiguity later in the paper, we will refer to the sensitive and resistant variants as ‘sub-strains’. Uninfected hosts (*U*) become infected with the resistant (*I_r_*) or the sensitive (*I_s_*) sub-strain at rate *β_r_* and *β_s_*; infections are cleared at rate *μ_r_* and *μ_s_*; the sensitive sub-strain experiences an additional clearance rate *τ* corresponding to the population antibiotic consumption rate (we assume immediate clearance following antibiotic exposure); and resistance is associated with a fitness cost affecting transmission (*β_r_* = *β_s_cβ*, where *c_β_* ≤ 1) and/or clearance (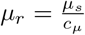, where *c_μ_* ≤ 1). The dynamics of this model are described by:

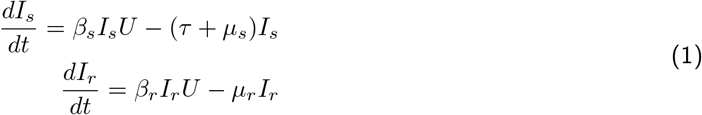

This system allows an equilibrium solution where both *I_s_* and *I_r_* are non-zero (i.e. stable coexistence of sensitivity and resistance) only when 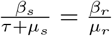. In other words, the resistant and sensitive sub-strains coexist only when their basic reproductive numbers (the average number of new infections an infected host gives rise to in a fully susceptible population) are equal. When this is not the case, the model predicts competitive exclusion: when resistance provides a fitness advantage 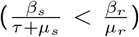, only the resistant sub-strain will be observed, and vice-versa when the sensitive sub-strain is fitter than the resistant sub-strain 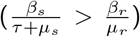. Defining overall fitness cost *c* = *c_μ_c_β_* and strain clearance rate *μ* = *μ_s_* to simplify notation, this threshold can be expressed as resistance being selected for when:

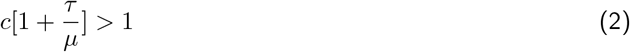

Thus, as reported previously [14], the fitness effect of resistance, which determines whether the resistant sub-strain out-competes the sensitive sub-strain, depends on the population antibiotic consumption rate, the fitness cost of resistance and the strain’s mean duration of carriage 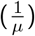, because longer carriage episodes have a greater risk of antibiotic exposure than shorter carriage episodes [14].

We now extend this model to *n* antibiotics: sub-strains can be either sensitive or resistant to each antibiotic, giving a total of 2*^n^* competing sub-strains. Similarly to the single drug model presented above, resistance to each antibiotic *j* has a transmission associated fitness cost *c_βj_* and/or a clearance associated fitness cost *c_μj_*. We assume no cost epistasis between resistance determinants: the fitness cost of resistance to an antibiotic does not depend on which other resistances are present on the sub-strain. Note that this assumption is not necessary for the demonstration of competitive exclusion in a multidrug context, but becomes important in later sections of paper. For consistency, we introduce it here. Therefore, a sub-strain *k*, resistant to the set of antibiotics *R*, has transmission rate *β_k_* = *β*Π_*j*∈*R*_ *c_βj_* and clearance rate 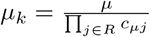 where *β* and *μ* are the transmission and clearance rates of the fully sensitive sub-strain. In addition, each sub-strain is cleared by the antibiotics it is sensitive to (set of antibiotics *S*), giving a total clearance rate of 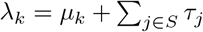, where *τ_j_* is the consumption rate of antibiotic *j*. The dynamics of each sub-strain are therefore described by:

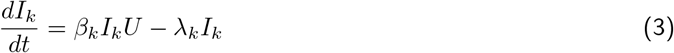

for all *k* ∈ {1, …, 2*^n^*} and with 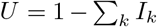. At equilibrium, 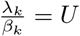 holds for all strains with non-zero frequency - we therefore recover the result from the single drug model: sub-strains can only coexist when they have the same reproductive number. When this is not the case, the frequency of resistance to each antibiotic is either 0% or 100% and a single resistance profile with the highest reproductive number 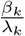 is expected to out-compete all others. In a multidrug context, therefore, coexistence-maintaining mechanisms are necessary to explain why multiple different resistance profiles are observed.

#### 2.1.2 Coexistence through heterogeneity in the fitness effect of resistance: single drug context

In this section, we note a structural similarity in plausible models of coexistence: a number of recently proposed coexistence mechanisms work by introducing variation in the fitness effect of resistance within either the host population or the pathogen population. We show that models with this structure can be simplified to a series of independent SIS models, which will allow us to gain insight into the pattern of association between resistance to different antibiotics in a multidrug context. We start by presenting a simple model for conceptual insights; additional complexity is explored in later sections.

In the first class of models we consider, coexistence arises from host population structure: assortatively mixing groups within the host population promote coexistence if the groups differ in the fitness effect of resistance, thus creating niches for resistance and sensitivity within the host population. This variation has been proposed to arise from between-group variation in the rate of antibiotic consumption (e.g. different antibiotic consumption in hospital and community settings [16], or between different age groups [15, 17]) or between-group variation in clearance rate (e.g. age-specific clearance rate [14]). For assortative mixing between host groups to promote coexistence, transmission between groups must be very low: even modest transmission between groups causes the groups to act as a single population and therefore abolishes coexistence [17] (see also Section 2.2). By treating this very low transmission as no transmission, we can represent host structure by modelling each of the host groups as a separate SIS model, with dynamics captured by Equation 1. In other words, we model competition between sensitivity and resistance within each host group as independent of the other host groups (Figure 1).

**Figure 1:**
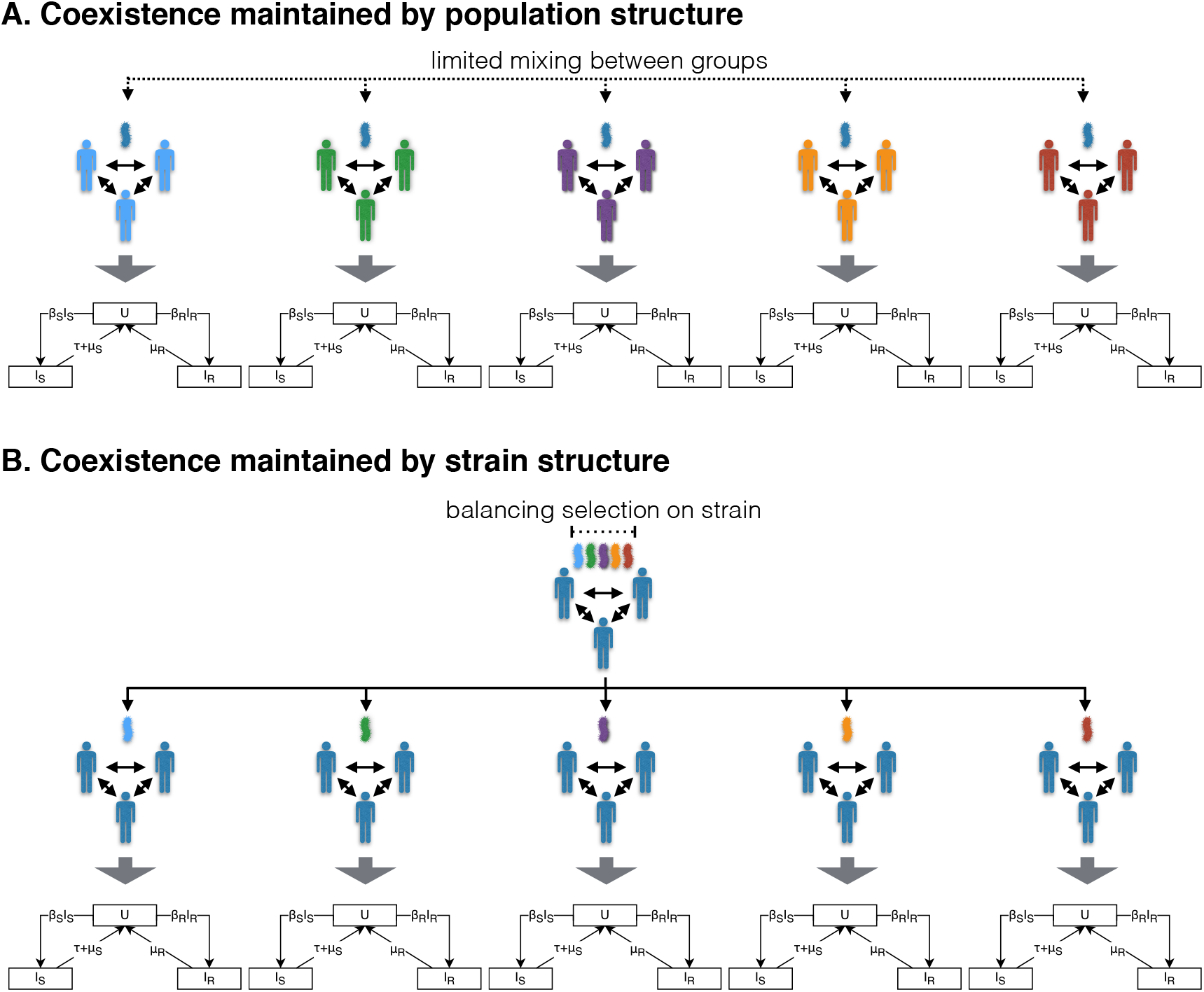
Illustration of how host population (panel A) and strain (panel B) structure maintain coexistence by introducing heterogeneity in the fitness effect of resistance and thus creating niches for sensitivity and resistance within the population. Each of the SIS model diagrams represents the resistance dynamics described by Equation 1. A: The resistance dynamics of assortatively mixing host groups can be modelled as independent SIS models by assuming no transmission between groups. Heterogeneity in the fitness effect of resistance arises from between host group differences in antibiotic consumption rate or clearance rate. B: The resistance dynamics of pathogen strains maintained by balancing selection can be modelled as independent SIS models by assuming no recombination. Heterogeneity in the fitness effect of resistance arises from between strain differences in clearance rate.

In the second class of models, coexistence arises from heterogeneity within the pathogen, rather than host, population: the heterogeneity in fitness effect of resistance arises from the presence of strains with different durations of carriage, maintained by balancing selection on the duration of carriage locus (e.g. serotype-specific acquired immunity allowing coexistence of serotypes with different durations of carriage in the pneumococcus [20]) [14]. Hence, similarly to assortatively mixing host groups in the first class of models, strains with different durations of carriage act as niches for sensitivity and resistance: coexistence is maintained by competition between sensitivity and resistance occurring independently within each strain. By assuming no recombination, we can again represent competition between sensitivity and resistance within each strain as a separate SIS model (Figure 1). Note that this class of models requires the presence of balancing selection maintaining diversity at the duration of carriage locus. In the simplified representation, this balancing selection is not modelled explicitly - coexistence of the strains differing in duration of carriage is assumed (see Supporting Information 1 for further discussion of this point).

Thus models in which coexistence arises from heterogeneity in the fitness effect of resistance - either within the host or pathogen population - can be represented by a series of independent SIS models. We refer to these individual SIS models as strata. In the case of a single strain circulating in a structured host population, the strata correspond to assortatively mixing host groups (e.g. age classes). In the case of multiple strains circulating in a homogeneous host population, the strata correspond to different strains (e.g. serotypes in the pneumococcus). When both strain and population structure are present, each stratum corresponds to a particular strain circulating in a particular host group. Following from Equation 2, resistance out-competes sensitivity in stratum *pi* (host group *p* and strain *i*) when:

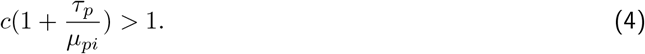

#### 2.1.3 Coexistence through heterogeneity in the fitness effect of resistance: multidrug context

We now extend this model to multiple antibiotics, which may differ in fitness cost and consumption rate. We assume no cost epistasis between resistance determinants: resistance to antibiotic *a* has the same fitness cost, *c_a_*, in presence and absence of resistance to antibiotic *b*. We also assume that different antibiotics are consumed in the same proportions in all host groups: antibiotic *a* accounts for proportion *γ_a_* of total antibiotic consumption, with 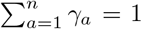, where *n* is the number of different antibiotics. The consumption rate of antibiotic *a* for host group *p* is therefore 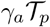, where 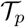 is the total antibiotic consumption rate of group *p*.

Under these assumptions, following from Equation 2, resistance to antibiotic *a* out-competes sensitivity in host group *p* and strain *i* when:

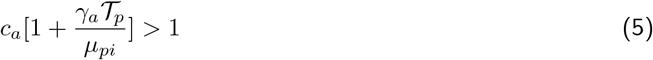

As before, there is no coexistence within the strata: below this threshold, sensitivity out-competes resistance. In a multi-drug context, a single resistance profile will therefore out-compete all others within each stratum.

Equation 5 simplifies the dynamics of multidrug resistance in that it does not account for the effect of exposure to antibiotics other than *a* or for the effect of the absence/presence of resistance to these antibiotics. In absence of resistance to other antibiotics, the clearance rate experienced by the strain depends on the rate at which these other antibiotics are consumed. In the presence of resistance to other antibiotics, the clearance rate may depend on the fitness cost of these resistances (if fitness cost affects clearance rate). We will pursue our reasoning using this simplified model because it captures the mechanism leading to the association between resistance to different antibiotics. The additional complexity arising from the effect of resistance on clearance rate does not meaningfully alter our results (Section 2.2).

#### 2.1.4 Heterogeneity in the fitness effect of resistance: predicted patterns of resistance

We can separate Inequality 5 into stratum (i.e host group and pathogen strain) and antibiotic related effects (left-hand and right-hand sides of Inequality 6, respectively):

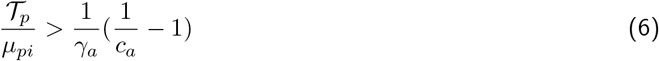

We call the ratio 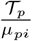 resistance proneness (*P_pi_*) and the ratio 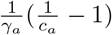 resistance threshold (*T_a_*). *P_pi_* reflects how advantageous resistance is within stratum *pi*. High antibiotic consumption (high *τ_p_*) and low clearance rate (low *μ_pi_*) lead to high resistance proneness. *T_a_* reflects how advantageous resistance against antibiotic *a* needs to be for it to be selected for. High fitness cost (low *c_a_*) and making up a low proportion of total antibiotic consumption (low *γ_a_*) lead to high resistance threshold. Rewriting Equation 6 using this notation, resistance to antibiotic *a* is selected for in stratum *pi* when:

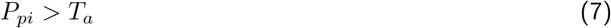

Resistance proneness depends only on stratum and resistance threshold depends only on antibiotic. As a consequence, the ordering of strata by resistance proneness is independent of antibiotic and the ordering of antibiotics by resistance threshold is independent of stratum. Therefore, for a set of *m* strata, with resistance proneness *P*_1_ < *P*_2_ < … < *P_m_* and for a set of *n* antibiotics, with resistance thresholds *T*_1_ < *T*_2_ < … < *T_n_*, the following will hold:

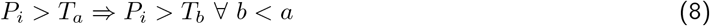

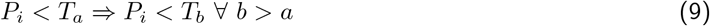

That is, if resistance to antibiotic *a* is selected, resistance to antibiotics with a lower resistance threshold will also be selected for. Conversely, if resistance to *a* is not selected for, resistance to antibiotics with a higher resistance threshold will also not be selected for. Thus, the ordering of antibiotics by resistance threshold (*T*_1_ < *T*_2_ < … < *T_n_*) determines the ordering of antibiotics by resistance frequency: the higher the resistance threshold, the lower the resistance frequency and resistance to a particular antibiotic will only be seen on resistance profiles with all more frequent resistances. Therefore, the fitness variation model predicts non-zero frequencies for only *n* + 1 out of the 2*^n^* possible resistance profiles: the only profile with resistance multiplicity of *m* (i.e. resistance to *m* antibiotics) will be the one with the *m* most common resistances (Figure 2). This pattern of resistance is referred to as ‘nested’ and predicts strong association between resistance to different antibiotics, with all resistance pairs in complete linkage disequilibrium (D’= 1, where D’ is the normalised coefficient of linkage disequilibrium (LD) - see Methods).

**Figure 2:**
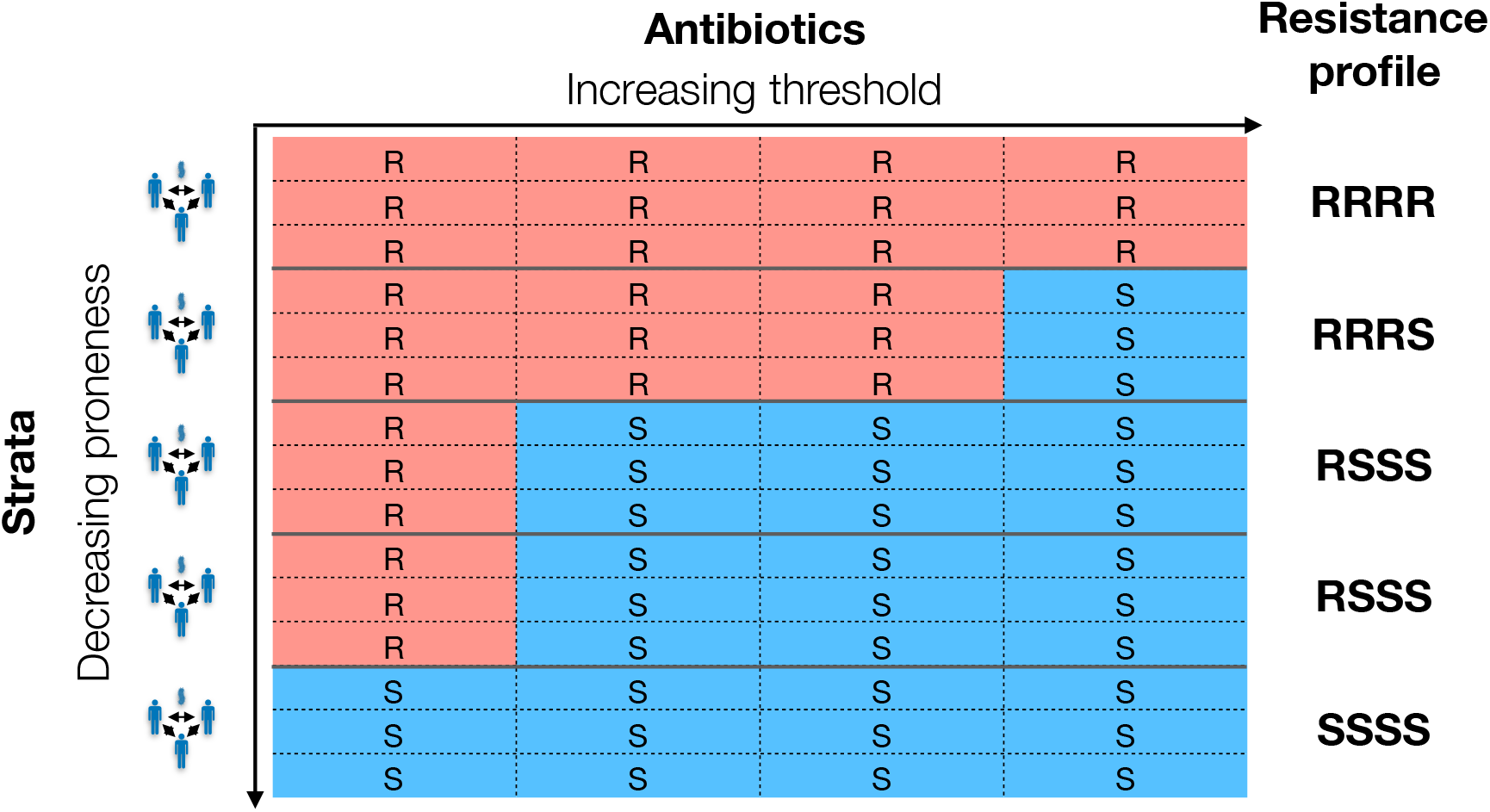
Example of a set of resistance profiles from a system with five strata and four antibiotics. Each row in the table corresponds to the resistance profile of one isolate - i.e. there are three isolates from each strata (equal sampling/size of strata is not necessary). Competitive exclusion within a stratum means all isolates from one stratum have the same resistance profile. The strata have been arranged from top to bottom in order of decreasing resistance proneness 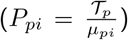. The antibiotics have been arranged left to right in order of increasing resistance threshold 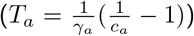, or, equivalently, decreasing resistance frequency. Resistance to a particular antibiotic outcompetes sensitivity in a stratum when the resistance proneness of the stratum is greater than the resistance threshold of the antibiotic. Resistance proneness being independent of antibiotic and resistance threshold being independent of stratum leads to nested resistance profiles (i.e. rarer resistances only observed in the presence of more common ones) and complete linkage disequilibrium between resistances. See Supporting Information 2 for an example of a set of non-nested resistance profiles.

### 2.2 Extension: additional complexity

In this section, we explore how introducing additional complexity to the simplified model affects our predictions about association between resistance determinants and nestedness.

#### 2.2.1 The effect of resistance on clearance rate

The model presented above ignores the effect of the absence/presence of resistance to other antibiotics on the resistance threshold of antibiotic *a*. As discussed, this is a simplification because i) in the absence of resistance against other antibiotics, the exposure to these antibiotics will contribute to clearance and ii) the presence of resistance to other antibiotics will affect clearance if the fitness cost of resistance increases clearance rate. In a multidrug context therefore, Equation 5 is an approximation.

In Supporting Information 3, we show that this approximation does not meaningfully affect our predictions about association between resistance determinants and nestedness of resistance profiles. The effect of resistance on clearance rate does not give rise to incomplete linkage disequilibrium (*D′* < 1) and non-nested resistance profiles, except under very specific circumstances: if the fitness cost of resistance affects clearance rate and more commonly prescribed antibiotics also have higher fitness cost. Even when incomplete linkage disequilibrium is possible theoretically, the parameter range under which it arises is extremely narrow (see Supporting Information 3), suggesting incomplete linkage disequilibrium and non-nested resistance profiles being observed because of the effect of resistance on clearance rate is unlikely.

#### 2.2.2 Intergroup transmission and recombination

In the model presented above, strata are fully independent, with no transmission between host groups and no recombination. As discussed, this is a simplification: coexistence can be maintained in the presence of mixing between strata if the rate of mixing is low enough [17]. In order to investigate the effect that mixing between strata has on our predictions about the association between resistance determinants, we construct i) a model with three different antibiotics and five host groups differing in clearance rate with transmission allowed between host groups and ii) a model with three different antibiotics and five strains differing in clearance rate with recombination allowed at the duration of carriage locus (see Methods for details). In both models, complete linkage disequilibrium is maintained in the presence of mixing between strata (Figure 3): mixing does not introduce any source of selection that would favour non-nested resistance profiles.

**Figure 3:**
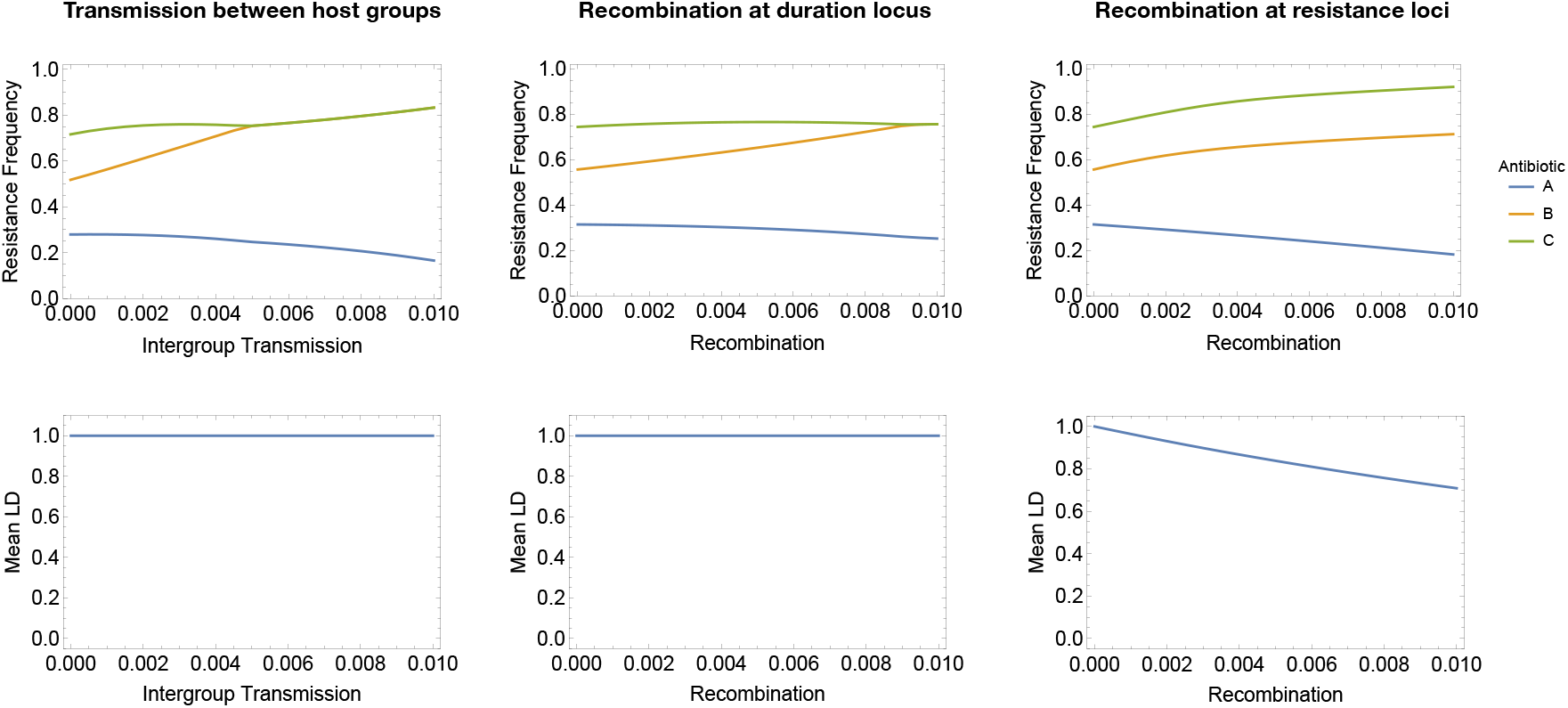
Strain frequencies and mean linkage disequilibrium (LD) between resistances in three models with three antibiotics (*A, B* and *C*) consumed at different rates. Left: a model with host population structure (five assortatively mixing host groups) with increasing levels of intergroup transmission. The rate of intergroup transmission on the x-axis (parameter *m* in the model represented by Equation 10, see Methods) reflects the proprotion of transmission events that occur between, instead of within, host group. Middle: a model with strain structure (five strains differing in duration of carriage) with increasing rates of recombination at the duration of carriage locus. Recombination rate on the x-axis (parameter *r* in the model represented by Equation 11, see Methods) reflects the probabilitity of co-infection, the probability of recombination occuring during co-infection and the probability of the recombinant strain being transmitted. Right: the same model with strain structure (five strains differing in duration of carriage) with increasing rates of recombination at the resistance loci.

We also investigate the effect of recombination at the resistance loci in the strain structured model (see Methods). (We do not implement recombination in the population structured model: we assume recombination requires co-infection and expect no co-infection in this model because of competitive exclusion within each host group.) Unlike recombination at the duration locus, recombination at the resistance loci breaks up linkage disequilibrium (Figure 3), decreasing the magnitude of association between resistance determinants. However, this effect is gradual and high levels of LD are maintained even at unrealistically high rates of recombination (see Supporting Information 4).

#### 2.2.3 Imperfectly correlated strata

In the simple model we present, the prediction of complete LD arises because we can separate the variation in the fitness effect of resistance into strata-related (i.e. pathogen and host) and antibiotic-related effects. This separability means that resistance proneness of a stratum is independent of antibiotic, which gives rise to complete LD between resistance to different antibiotics and resistance profiles with nested structure. This separability requires two assumptions: first, that the fitness cost of a particular resistance is the same in all strata (i.e. no variation in the fitness cost of resistance between strains) and second, that different types of antibiotics are consumed in the same proportions in all strata (i.e. variation in the *rate* at which host groups consume antibiotics, but not in the mixture of antibiotic types).

Both of these requirements may represent oversimplifications of resistance dynamics. In the case of the first assumption, there is no direct evidence for stable variation in the fitness cost of resistance (i.e. variation in fitness cost maintained by balancing selection, preventing the lower fitness cost phenotype reaching fixation). However, the processes determining fitness cost are not fully understood and the fitness cost of resistance mutations is thought to depend on both genetic background and environment [21]. It is therefore difficult to rule out that stable between-strain variation might exist. Similarly, the assumption about antibiotic prescription rates is generally plausible, but might not hold in all contexts. For example, prescription rates of different antibiotic classes are highly correlated between US States (mean correlation over 0.9, based on data from the Center for Disease Dynamics, Economics & Policy, available at https://resistancemap.cddep.org). However, correlations between relative prescription rates may not be so high between all host groups, such as children and adults. For example, fluoroquinolones are primarily used in adults but not children [22] which may explain why association between resistances is weaker for fluoroquinolones than for other antibiotics [2].

To test the extent to which antibiotic-specific resistance proneness affects predictions about resistance profiles, we simulate strata with increasingly uncorrelated resistance proneness for different antibiotics (see Methods). The frequency of nested resistance profiles and the average linkage disequilibrium decreased with decreasing correlation (Figure 4). However, the effect was gradual: imperfectly correlated resistance proneness gives rise to relatively high frequencies of nested resistance profiles and above zero mean *D′*. The association between resistance determinants is only lost (mean *D′* = 0) when resistance proneness is fully uncorrelated. The fitness variation model therefore predicts association between resistance determinants even when the fitness variation giving rise to coexistence is not identical for all antibiotics.

**Figure 4:**
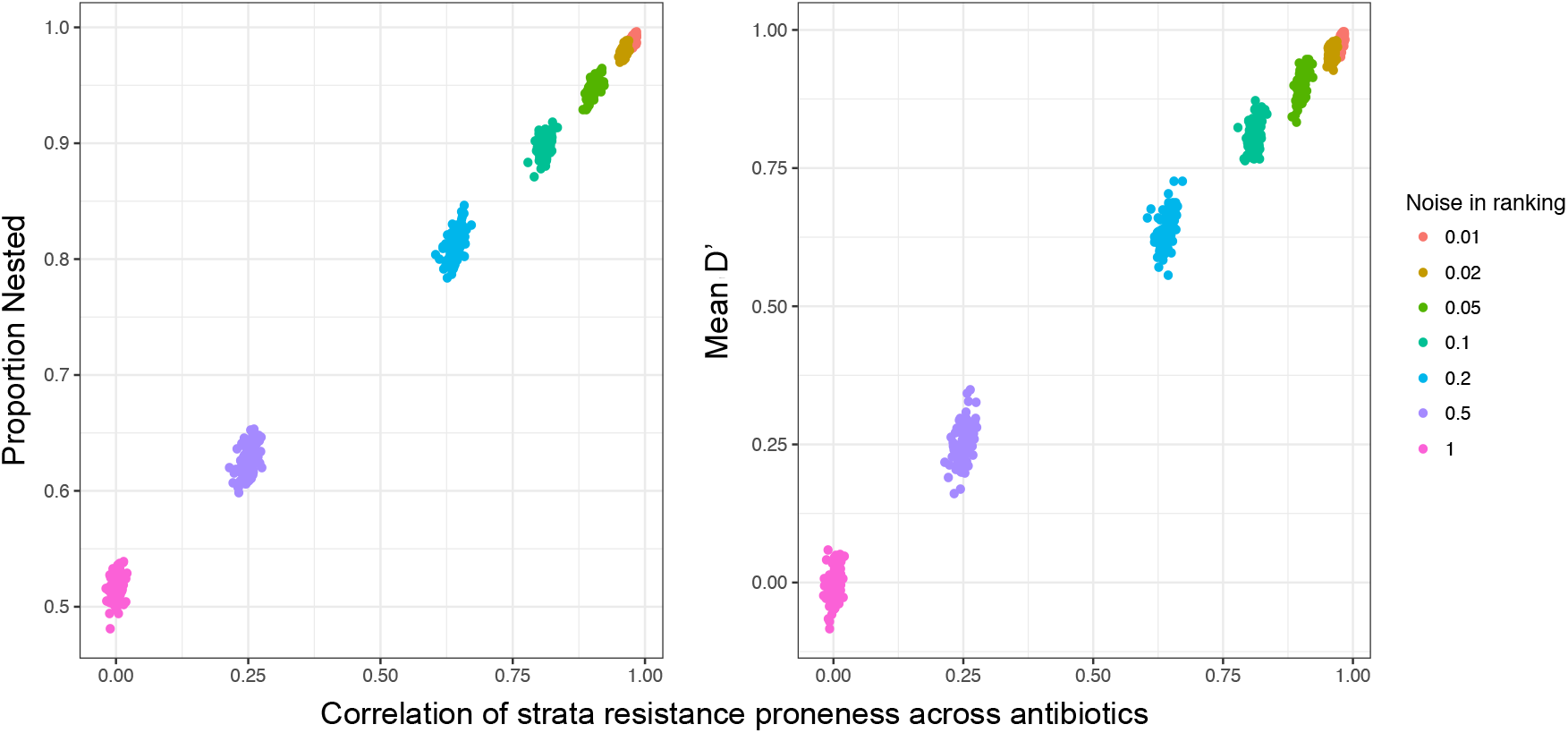
The extent of predicted association between resistance to different antibiotics depends on the extent to which the fitness effect of resistance is correlated for the different antibiotics. The figure shows the proportion of nested resistance profiles and mean *D′* between resistance pairs for simulated data with varying concordance in resistance proneness to different antibiotics (see Supporting Information 2 for examples of nested and non-nested resistance profiles). Concordance in resistance proneness is measured as spearman correlation in resistance proneness across antibiotics. Each data point corresponds to one simulated dataset (1000 simulated datasets for each level of discordance, see Methods). The simulated datasets have the same dimensions and resistance frequencies as the Maela pneumococcal dataset (Table 2). The proportion of nested resistance profiles in the Maela dataset is 0.71 and the mean *D′* is 0.46.

### 2.3 Model predictions are consistent with trends in bacterial datasets

#### 2.3.1 High levels of association between resistance determinants

The fitness variation model predicts high levels of linkage disequilibrium between resistance to different antibiotics and a high proportion of resistance profiles with a nested structure. We measured these quantities in six bacterial datasets for which data on resistance to multiple antibiotics was available (four hospital datasets from the United States for different species and two pneumococcal datasets from Massachusetts and Maela - see Methods for additional details) and found trends consistent with our predictions (Table 2).

**Table 2:**
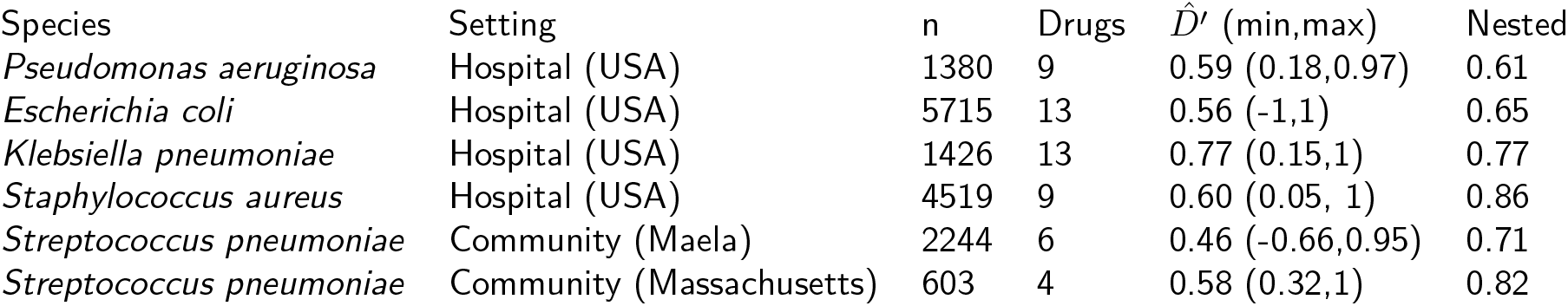
Mean pairwise LD between antibiotic pairs 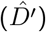 and proportion of resistance profiles that are nested for six bacterial datasets (see Methods for details of datasets). The “n” column indicates the number of isolates in the dataset and the “drugs” column indicates the number of drugs resistance was tested for. Positive *D′* indicates resisistance determinants tend to appear together. The values in parentheses give the range of pairwise *D′* in the dataset. Note that in the instances where the minimum *D′* value is negative or close to zero, at least one of the resistances in the pair this *D′* value corresponds to is present at very low frequency in the dataset (0.08% for the *E. coli* dataset, 1.7% in the *S. auerus* dataset, 6.1% for the Maela pneumococcal dataset).

| Species | Setting | n | Drugs | $\hat{D}'$ (min,max) | Nested |
| --- | --- | --- | --- | --- | --- |
| *Pseudomonas aeruginosa* | Hospital (USA) | 1380 | 9 | 0.59 (0.18,0.97) | 0.61 |
| *Escherichia coli* | Hospital (USA) | 5715 | 13 | 0.56 (-1,1) | 0.65 |
| *Klebsiella pneumoniae* | Hospital (USA) | 1426 | 13 | 0.77 (0.15,1) | 0.77 |
| *Staphylococcus aureus* | Hospital (USA) | 4519 | 9 | 0.60 (0.05, 1) | 0.86 |
| *Streptococcus pneumoniae* | Community (Maela) | 2244 | 6 | 0.46 (-0.66,0.95) | 0.71 |
| *Streptococcus pneumoniae* | Community (Massachusetts) | 603 | 4 | 0.58 (0.32,1) | 0.82 |

#### 2.3.2 Duration of carriage predicts resistance multiplicity

The fitness variation model predicts that duration of carriage and antibiotic consumption rate within strata will determine resistance multiplicity. Fully testing this prediction is challenging, because we do not have a full understanding of which host and pathogen characteristics are relevant in defining the strata. To partially test the prediction, we test the association between duration of carriage and resistance using a dataset of pneumococcal carriage episodes and associated durations of carriage [23] (‘Maela dataset’, see Methods). The average resistance multiplicity of a serotype is indeed positively associated with the serotype’s average duration of carriage (Kendall rank correlation 0.27 95% CI 0.05-0.46, n = 38 excluding serotypes with fewer than 10 observations). A caveat here is that the direction of causality for this association is not entirely clear: we suggest long duration of carriage selects for resistance but resistance would also be expected to lead to longer duration of carriage through decreased clearance from antibiotic exposure. However, at the serotype level, differences in duration of carriage are thought to arise from the properties of serotype capsules [24] (rather than differences in antibiotic resistance). This suggests that longer duration of carriage favouring resistance does indeed contribute to the association between duration of carriage and resistance multiplicity.

## 3 Discussion

### 3.1 Generalised model of coexistence predicts high frequencies of multidrug resistance

In this paper, we approach the question of explaining observed patterns of association between resistance to different antibiotics (‘MDR over-representation’) in terms of understanding the competition between strains with different resistance profiles. We consider recent models of coexistence [16, 15, 14, 17] in which coexistence is maintained by heterogeneity in the fitness effect of resistance, arising either from heterogeneity in the rate of antibiotic consumption and/or difference in duration of carriage. We present a generalised version of these types of models, in which competition between antibiotic sensitivity and resistance is simplified to a series of independent sub-models (strata). We show that this model structure also gives rise to MDR over-representation because resistance to all antibiotics will be selected for in the strata where the fitness benefit of resistance (‘resistance proneness’) is the highest. Therefore, our results suggest that two pervasive trends in resistance dynamics, the robust coexistence of antibiotic sensitive and resistant strains and the over-representation of multidrug resistance, can both be explained by heterogeneity in the fitness effect of resistance in the pathogen population.

We first present a simplified model for conceptual insights and then explore how additional complexity affects predicted trends. Under the strong assumption of identical antibiotic prescription patterns in all strata and no recombination, this model predicts complete linkage disequilibrium (*D′* = 1) between resistance to all antibiotics. Relaxing these assumption decreases the magnitude of linkage disequilibrium, giving rise to values of *D′* similar to those observed in multiple bacterial datasets. High *D′* is maintained even at unrealistically high recombination rates. The association between resistance determinants is only abolished when resistance proneness between strata is completely uncorrelated. Thus, even in context where patterns of prescription differ considerably between host groups, we would still expect a degree of association between resistance determinants when variation in duration of carriage contributes to variation in the fitness effect of resistance.

Our results therefore show that when variation in the fitness effect of resistance is present and when this variation is at least partially correlated for different antibiotics, it will give rise to MDR over-representation. The extent to which this mechanism accounts for observed patterns of MDR over-representation therefore depends on the extent to which this type of fitness variation is present in pathogen populations.

It is not entirely straightforward to evaluate how common variation in the fitness effect of resistance is. Wide-spread coexistence of sensitivity and resistance is not direct evidence for the pervasiveness of fitness variation because coexistence may not always arise through this mechanism. Although the majority of mechanisms proposed to date [16, 15, 14, 17] work through fitness variation, other mechanisms are also possible [13]. In particular, recent modelling suggests that co-infection with sensitive and resistant strains gives rise to frequency-dependent selection for resistance and thus promotes coexistence [18]. However the magnitude of this effect depends on the nature of within-host competition [18], which there is limited data about. Thus while theoretically plausible, the extent to which this mechanism contributes in practice is still unclear. It is worth noting that different coexistence mechanisms are not mutually exclusive. If coexistence arises through a combination of fitness variation and other mechanisms, we would *a priori* still expect the fitness variation to give rise to MDR over-representation.

In the work presented here, we consider fitness variation arising from heterogeneity in antibiotic consumption between host groups (hospitals vs communities, geographic regions, age classes) and from heterogeneity in duration of carriage between host groups (age classes) and between strains (pneumococcal serotypes). This is not an exhaustive list of possible sources of heterogeneity. For example, serotype does not fully account for heritable variation in pneumococcal duration of carriage [23], suggesting other genetic traits also play a role in determining carriage duration. In light of recent results suggesting wide-spread negative frequency-dependent selection in bacterial genomes [25, 26], it is not implausible to suggest these duration of carriage loci may also be under frequency-dependent selection. If so, diversity at these loci would create another source of variation in the fitness effect of resistance and hence promote coexistence and MDR over-representation. More broadly, variation in the fitness effect of resistance may arise through different mechanisms for pathogens with a different ecology than modelled in this work. For example, we have modelled a pathogen that is mostly carried asymptomatically and therefore exposed primarily to antibiotics prescribed against other infections. For pathogens where antibiotics prescribed due to infection with the pathogen itself contribute to a significant proportion of antibiotic exposure, the presence of strains differing in invasiveness would give rise to between-strain variation in antibiotic exposure and heterogeneity in the fitness effect of resistance.

In summary, fitness variation is likely contributes to the pervasive coexistence of antibiotic sensitivity and resistance and therefore plays a role in wide-spread MDR over-representation, although this does not preclude a potential role for other mechanisms in contributing to the trend (Table 1).

Finally, although the model builds on work exploring the stable coexistence of antibiotic sensitivity and resistance and coexistence is robustly observed in multiple datasets, the prediction that variation in the fitness effect of resistance leads to MDR over-representation does not require coexistence to be stable. We would expect MDR over-representation in the presence of fitness variation, even when this variation is not enough to maintain stable coexistence: for all antibiotics, the increase of resistance frequencies towards fixation would occur most rapidly in the populations with the greatest selection pressure for resistance. Under these circumstances, fitness variation would give rise to transient MDR over-representation.

### 3.2 Public health implications

From a public health perspective, the fitness variation model makes two concerning predictions. Firstly, we predict frequencies of pan-resistance will be high: in a perfectly nested set of resistance profiles, the frequency of pan-resistance is equal to the frequency of the rarest resistance. As a consequence, we would expect resistance arising in response to adoption of new antibiotics or increased usage of existing antibiotics to appear on already multidrug resistant lineages - an observation which has been made for the emergence of ciprofloxacin resistance in *N. gonorrhoeae* in the United States [27].

Secondly, our analysis has implications for the effectiveness of potential interventions against MDR. The variation in the fitness effect of resistance to different antibiotics need not be perfectly correlated for it to promote MDR over-representation. If the variation in fitness effect is maintained by multiple factors (e.g. differential antibiotic consumption between populations and variation in clearance rates), removing one of these factors (e.g. changing patterns of prescription) may have limited impact on MDR.

The fitness variation model provides an explanation for MDR over-representation that is consistent with long term stability in resistance frequencies. This is relevant when considering temporal trends in resistance frequencies and predicting the future burden of resistance: other explanations for MDR over-representation (e.g. cost epistasis, correlated antibiotic exposure at the individual level - see Table 1) often require MDR strains to have an overall fitness advantage over strains with lower resistance multiplicity. This would imply that the higher than expected frequency of MDR is evidence for MDR strains out-competing other strains and thus suggest that MDR strains will eventually take over. Conversely, in the model we present, MDR strains are not out-competing other strains: all resistance frequencies are at equilibrium and MDR over-representation arises from the distribution of resistance determinants. It is worth noting, however, that even in the context of the fitness variation model, on a very long time-scale, we might expect the frequency of resistance to rise if bacteria are able to evolve resistance mechanisms that carry a lower fitness cost.

### 3.3 Conclusion

We show that previously proposed models in which coexistence of antibiotic sensitivity and resistance is maintained by heterogeneity in the fitness effect of resistance also predict high frequencies of multidrug resistance. The pervasive trends of coexistence and MDR over-representation can therefore be considered, at least partially, facets of the same phenomenon. We do not propose that the model we present fully explains observed patterns of association between resistance determinants. However, this effect should be considered when evaluating the role of antibiotic-specific MDR promoting mechanisms. From a public health point of view, the model we present is concerning because it predicts high frequencies of pan-resistance. On the other hand, heterogeneity in the fitness effect of resistance as an explanation for MDR over-representation allows reconciling this trend with long term stability in resistance frequencies.

## 4 Methods

### 4.1 Datasets

The Maela pneumococcal dataset [28], collected from a refugee camp on the border of Thailand and Myanmar from 2007 to 2010, consisted of 2244 episodes of carriage, with associated antibiograms and carriage durations. Data were obtained from, and durations of carriage calculated by, Lees et al. [23] (Supporting Data 1). Data on antibiotic sensitivity was provided for ceftriaxone, chloramphenicol clindamycin, erythromycin, penicillin, co-trimoxazole (trimethoprim/sulfamethoxazole) and tetracycline. Ceftriaxone was excluded from the analysis because data was missing for a large proportion of isolates (44%). The Massachusetts pneumococcal dataset, collected as part of the SPARC (*Streptococcus pneumoniae* Antimicrobial Resistance in Children) project [29], was obtained from Croucher et al. (2013) [30] (data available from Croucher et al [30]). Croucher et al. reported minimum inhibitory concentrations (MICs) for penicillin, ceftriaxone, trimethprim, erithromycin, tetracycline and chloramphenicol. Tetracycline and chloramphenicol were excluded from the analysis because data was missing for a large proportion of isolates (47% and 67% respectively). Non-sensitivity was defined in accordance to pre-2008 Clinical and Laboratory Standards Institute breakpoints [31]. For both datasets, ‘resistance’ as used throughout the paper refers to non-sensitivity. The four hospital datasets were obtained from Chang et al. [2] (Supporting Data 2).

### 4.2 Linkage disequilibrium

If the frequency of resistance to antibiotic *a* is *p_a_* and the frequency of resistance to antibiotic *b* is *p_b_*, the coefficient of linkage disequilibrium between resistance to antibiotics *a* and *b* is *D_ab_* = *p_ab_* − *p_a_p_b_*, where *p_ab_* is the frequency of resistance to both *a* and *b*. The normalised coefficient *D′_ab_* is given by: 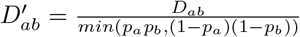 if *D_ab_* < 0 and 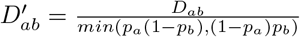 if *D_ab_* > 0.

In general the sign of *D^′^* is arbitrary because it depends on which alleles are chosen for the calculation. We consistently calculate *D′* using the frequency of resistance: positive D′ therefore means resistance to one antibiotic is associated with resistance to the other, while negative *D′* means association between sensitivity and resistance.

We use *D′* as a measure of linkage disequilibrium (as opposed to the other commonly used metric *r*^2^) because the fitness effect model makes prediction specifically about *D′*. The coefficient of determination, *r*^2^, measures the extent to which isolates that are resistant to one antibiotic are also resistant to another antibiotic while *D′* captures the extent to which resistance to two antibiotics will be found in the same isolates, given the observed resistance frequencies. Therefore, *r*^2^ is affected by how similar resistance frequencies are and by the distribution of the resistant determinants, while *D′* is only affected by the latter.

### 4.3 Effect of intergroup transmission and recombination

To test the effect of relaxing the assumption that the pathogen dynamics can be divided into non-interacting sub-models, we include three additional models.

First, we model the dynamics of resistance to three antibiotics (i.e. eight possible resistance profiles) spreading in a host population consisting of five host groups. The antibiotics make up different proportions of total antibiotic consumption (20, 35 and 45% of total antibiotic consumption rate *τ*). The pathogen experiences a different clearance rate within each host class *p* (*μ_p_*). In addition, sub-strain with resistance profile *g* experiences clearance from antibiotic exposure at rate *τ_g_* which depends on its resistance status: *τ_g_* = *τ*(*i_a_*0.20 + *i_b_*0.35 + *i_c_*0.45), where *i_a_* = 1 if *g* is sensitive to antibiotic *a* and 0 otherwise. Resistance to each antibiotic decreases transmission rate by a factor of *c*. Uninfected hosts of class *p* (*U_p_*) are therefore infected at rate 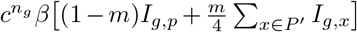, where *n_g_* is the number of antibiotics strain *g* is resistant to, *m* is a parameter that sets the extent of mixing between the classes and *P′* is the set of population classes excluding *p*. The dynamics of strain *g*within population *p* are thus described by:

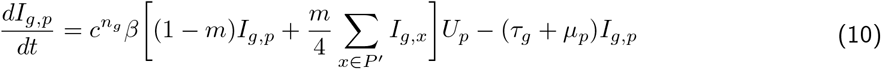

Second, we model the dynamics of resistance to three antibiotics in a single host population in pathogen with five strains differing in clearance rate (i.e. eight resistance profiles and five strains, giving a total of 40 possible sub-strains) with recombination at the duration of carriage locus. Strain *i* is cleared at rate *μ_i_* and, as above, sub-strains with resistance profile *g* experience clearance from antibiotic exposure at rate *τ_g_* which depends on its resistance status: *τ_g_* = *τ*(*i_a_*0.20 + *i_b_*0.35 + *i_c_*0.45). Resistance to each antibiotic decreases transmission rate by a factor of *c*. Balancing selection is modelled similarly to Lehtinen et al. [14], by scaling transmission rate of strain i by a factor *ψ_i_* which depends on the strain’s prevalence: 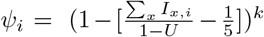, where *k* is a parameter setting the strength of balancing selection and *U* is the uninfected host class. Recombination at the duration of carriage locus is modelled by allowing hosts infected with strain i with resistance profile *g* to transmit strain *j* with resistance profile *g* at a rate 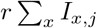. Recombination therefore decreases the transmission of strain i with resistance profile *g* by 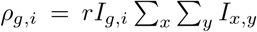 and increases it by 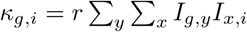. Note that the recombination rate parameter *r* captures the probability of co-infection, the probability of recombination occurring and the probability of transmitting the recombinant sub-strain. The dynamics of strain *i* with resistance profile *g* are described by:

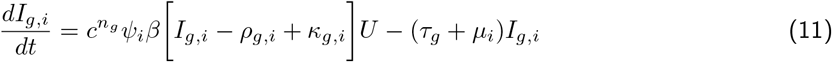

The third model is the same as the one above, with the exception that recombination occurs at the resistance loci instead of the duration of carriage locus. It is therefore described by Equation 11, but the expressions for *ρ* and *κ* are different. We define resistance profile 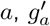 as a resistance profile otherwise identical to *g*, but with the other allele at locus *a* (i.e. if *g* is sensitive to antibiotic 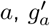 is resistant), *N_g,a_* as the set of resistance profiles with the same allele at locus *a* as profile *g* and 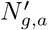 as the set of resistance profiles with the different allele at locus *a* than profile *g*. Hosts infected with strain *i* with resistance profile *g* transmit a strain *i* with a resistance profile 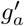 at rate 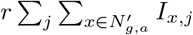. Recombination can occur at any of the three resistance loci (we assume recombination rates are low enough to ignore the possibility of recombination occurring at multiple loci at the same time). Recombination therefore decreases the transmission of strain *i* with resistance profile *g* by 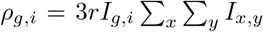 and increases it by 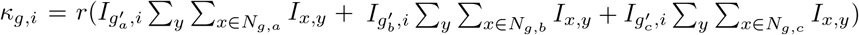.

The parameter values for the results presented in Figure 3 are: *c* = 0.95, *β* = 2, {*μ*_1_, .., *μ*_5_} = {1.2, 1., 0.8, 0.6, 0.4}, *τ* = 0.12 and *k* = 5.

### 4.4 Effect of imperfectly correlated strata resistance proneness

To test the effect of relaxing the assumptions that lead to strata resistance proneness being identical for all antibiotics (i.e. antibiotics consumed in the same proportions within all strata and the cost of resistance being the same within all strata), we created simulated resistance data similar to the Maela dataset (2244 isolates, 6 antibiotics). We randomly assigned a resistance proneness (i.e. 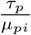) to each isolate. We then set resistance thresholds for each antibiotic. Isolates with a resistance proneness above this threshold were resistant. (The resistance thresholds were chosen to give the same resistance frequencies as observed in the Maela data). We then calculated the proportion of nested resistance profiles for this simulated dataset. To generate imperfectly correlated proneness, we replicated the resistance proneness vector for each antibiotic to generate a resistance proneness matrix. In this initial matrix, each isolate had identical resistance proneness for all antibiotics. We then redrew resistance proneness values for a proportion of randomly chosen entries in this matrix (proportion labelled as ‘noise in rankings’in Figure 4), thus generating differing resistance proneness to different antibiotics. Resistance was then assigned and nestedness calculated as above. By increasing the proportion of redrawn entries, we generated increasingly uncorrelated fitness advantage rankings.

