## Supporting Information for "On the evolutionary ecology of multidrug resistance in bacteria"

### On the evolutionary ecology of multidrug resistance in bacteria: Supporting Information

#### 1 Implicit modelling of balancing selection

In the main text, we show that for the purposes of understanding association between resistance determinants, models of competition between antibiotic resistance and sensitivity with host population structure and strain structure can both be simplified to a series of independent SIS models. In the case of strain structure, this requires the presence of balancing selection to maintain the strains differing in duration of carriage. We do not model this balancing selection explicitly: we assume coexistence of the strains and focus on modelling the competition between sensitivity and resistance within each strain.

To illustrate this simplification, we consider a model with $n$ strains in which strain coexistence is maintained by scaling the transmission rate of strain $i$ ($\beta_i$) by $g(f_i)$, where $f_i$ is the frequency of strain $i$ and $g()$ is a decreasing function (this is the model in Lehtinen et al. [1]). For any strain $i$, with sensitive and resistant sub-strains $is$ and $ir$, clearance rate $\mu_i$, cost of resistance $c_\beta$ ($c_\beta \leq 1$) and $c_\mu$ ($c_\mu \leq 1$) and population antibiotic consumption $\tau$, the competition between sensitivity and resistance is captured by:

$$\begin{aligned} \frac{dI_{is}}{dt} &= \beta_i I_{is} g(f_i) U - (\tau + \mu_i) I_{is} \\ \frac{dI_{ir}}{dt} &= c_\beta \beta_i I_{ir} g(f_i) U - \frac{\mu_i}{c_\mu} I_{ir} \end{aligned} \tag{1}$$

This model predicts competitive exclusion within strain: when $c_\beta c_\mu(1 + \frac{\tau}{\mu_i}) > 1$, strain $i$ will be resistant ($I_{is} = 0$, $I_{ir} > 0$) and when $c_\beta c_\mu(1 + \frac{\tau}{\mu_i}) < 1$, strain $i$ will be sensitive ($I_{is} > 0$, $I_{ir} = 0$). Importantly, these expression depend only on strain $i$: the competition between sensitivity and resistance within strain $i$ is independent of the other strains in the model. For our purposes, therefore, we can model each strain separately, replacing $g(f_i)U$ by a fixed number of hosts available to strain $i$, giving rise to the model presented in the main text.

#### 2 Schematics of resistance profiles

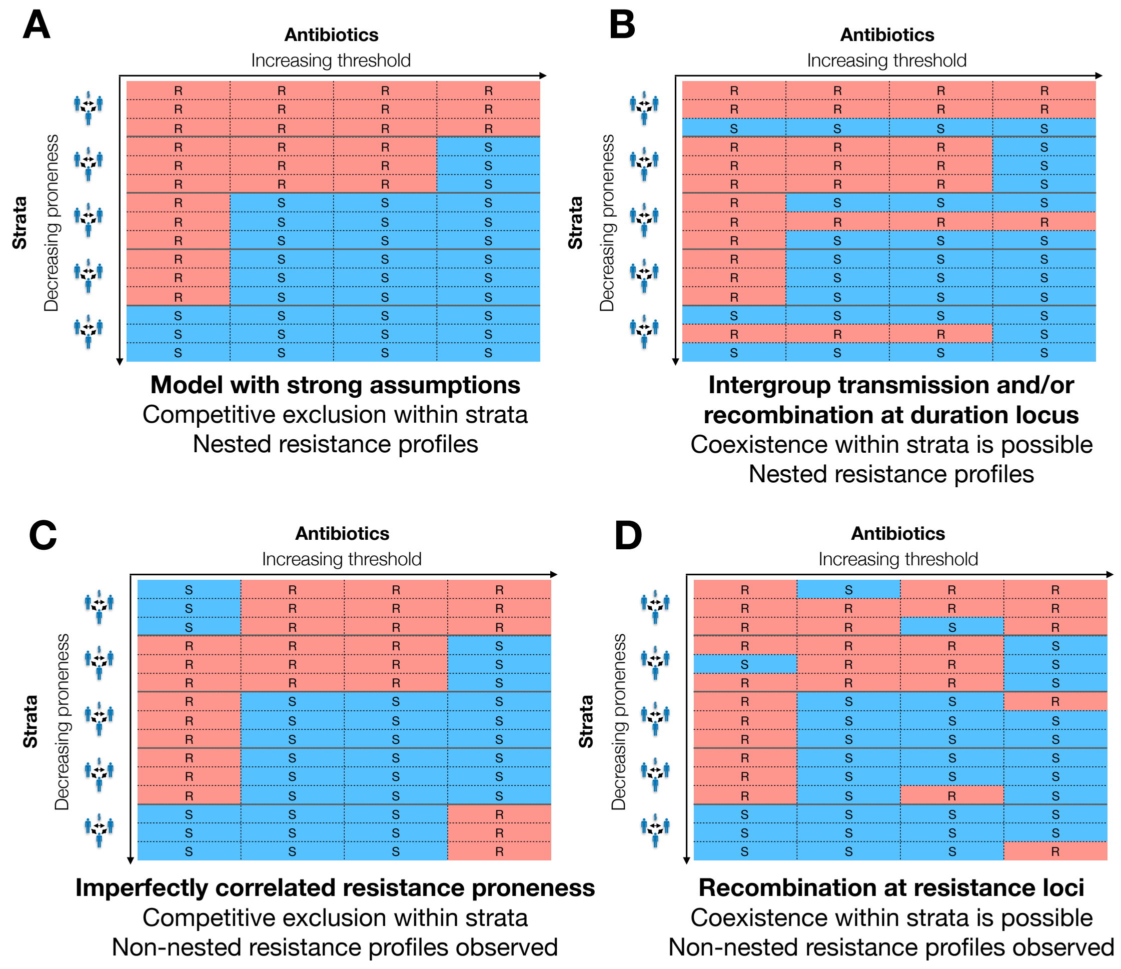


Figure 1: Schematic illustrating the effect of relaxing various model assumption on a set of resistance profiles from a system with five strata and four antibiotics. A: set of resistance profiles under the model with strong assumptions (main text Figure 2). B: Mixing between strata (integroup transmission or recomination at the duration locus) introduces some within-strata diversity in resistance profiles, but all observed resistance profiles have a nested structure. C: If the resistance proneness of a strata is not the same for all antibiotics, non-nested resistance profiles can arise. D: Recombination at the resistance loci breaks down associations between resistances and leads to the presence of non-nested resistance profiles and coexistence of multiple resistance profiles within strata.

#### 3 Effect of resistance on clearance rate in multidrug model

In Sections 2.1.3 and 2.1.4 of the main text, when considering selection for resistance against antibiotic $a$, we formulate the model in a way that ignores the effect of treatment with, and resistance against, *other* antibiotics. Resistance against other antibiotics affects clearance rate: firstly, by determining whether the strain can be cleared through exposure to the other antibiotics and, secondly, through the effect of fitness cost of resistance (if applicable). Therefore, the absence/presence of resistance against other antibiotics will affect selection for resistance to antibiotic $a$.

To investigate the impact this effect has on our predictions about the association between resistance determinants, we consider a system with two antibiotics and competition between four resistance profiles: sensitive to both antibiotics ($SS$); sensitive to antibiotic 1, resistant to antibiotic 2 ($SR$); resistant to antibiotic 1, sensitive to antibiotic 2 ($RS$); resistant to both ($RR$). Within a particular stratum with total antibiotic consumption $\tau$ and clearance rate $\mu$, the dynamics of these strains are captured by:

$$\begin{aligned}
\frac{dI_{SS}}{dt} &= \beta I_{SS}U - (\tau + \mu)I_{SS} \\
\frac{dI_{SR}}{dt} &= c_{\beta 2}\beta I_{SR}U - (\gamma_1\tau + \frac{\mu}{c_{\mu 2}})I_{SR} \\
\frac{dI_{RS}}{dt} &= c_{\beta 1}\beta I_{RS}U - (\gamma_2\tau + \frac{\mu}{c_{\mu 1}})I_{RS} \\
\frac{dI_{RR}}{dt} &= c_{\beta 1}c_{\beta 2}\beta I_{RR}U - \frac{\mu}{c_{\mu 1}c_{\mu 2}}I_{RR}
\end{aligned} \tag{2}$$

where $U + I_{SS} + I_{SR} + I_{RS} + I_{RR} = 1$; $\beta$ is the transmission rate; $c_{\beta 1} \leq 1$ and $c_{\beta 2} \leq 1$ are the cost of resistance to antibiotic 1 and 2 respectively on transmission and $c_{\mu 1} \leq 1$ and $c_{\mu 2} \leq 1$ are the costs on clearance; and $\gamma_1$ and $\gamma_2$ are the proportion of total antibiotic consumption corresponding to each antibiotic. Without loss of generality, we assume antibiotic 2 is prescribed at a higher rate than antibiotic 1 ($0.5 > \gamma_1 = 1 - \gamma_2$).

This system will give rise to incomplete linkage disequilibrium ($D' < 1$) between the resistances (and non-nested resistance profiles) if there are strata (i.e. some values of resistance proneness $\frac{\tau}{\mu}$) where strain $RS$ is selected for and other strata (i.e. other values of $\frac{\tau}{\mu}$) where strain $SR$ is selected for. The basic reproductive number ($R0$) of $RS$ is $\frac{c_{\beta 1}\beta}{\frac{\mu}{c_{\mu 1}} + \gamma_2\tau}$ and the basic reproductive number of $SR$ is $\frac{c_{\beta 2}\beta}{\frac{\mu}{c_{\mu 2}} + \gamma_1\tau}$. $SR$ out-competes $RS$ when $R0_{SR} > R0_{RS}$. Re-arranging this gives:

$$\frac{\tau}{\mu} > \frac{c_{\beta 1}c_{\mu 1} - c_{\beta 2}c_{\mu 2}}{c_{\mu 1}c_{\mu 2}(\gamma_2 c_{\beta 2} - \gamma_1 c_{\beta 1})}$$

When the fitness cost of the more commonly prescribed antibiotic (antibiotic 2) is lower ($c_{\beta 1}c_{\mu 1} < c_{\beta 2}c_{\mu 2}$, $c_{\beta 1} \leq c_{\beta 2}$ and $c_{\mu 1} \leq c_{\mu 2}$), the inequality will hold independent of the value of $\mu$ and $\tau$ (as long as $u > 0$ and $\tau \geq 0$): the left-hand side is always positive, whereas the right-hand side is always negative. Thus, resistance to the more commonly prescribed antibiotic with lower fitness cost is always more beneficial than resistance to the other drug and $SR$ is fitter than $RS$ in all strata.

When the fitness cost of the more commonly prescribed antibiotics is greater ($c_{\beta 1}c_{\mu 1} > c_{\beta 2}c_{\mu 2}$, $c_{\beta 1} \geq c_{\beta 2}$ and $c_{\mu 1} \geq c_{\mu 2}$), the relative fitness of $SR$ and $RS$ depends on the antibiotic consumption rate and clearance rate and may therefore differ between strata. However, incomplete linkage disequilibrium will only arise if a change in $\frac{\tau}{\mu}$ leads from a stable equilibrium with $SR$ at fixation to a stable equilibrium with $RS$ at fixation (or, equivalently, vice versa).

We perform a stability analysis to determine the parameter space in which this transition occurs. In line with the result above, for the $SR$ equilibrium to be stable requires:

$$\frac{\tau}{\mu} > \frac{c_{\beta 1}c_{\mu 1} - c_{\beta 2}c_{\mu 2}}{c_{\mu 1}c_{\mu 2}(\gamma_2 c_{\beta 2} - \gamma_1 c_{\beta 1})}$$

In addition, the following conditions must also be met:

$$\frac{\tau}{\mu} < \frac{1 - c_{\mu 1}c_{\beta 1}}{\gamma_1 c_{\mu 1}c_{\mu 2}c_{\beta 1}}$$
$$\frac{\tau}{\mu} > \frac{1 - c_{\mu 2}c_{\beta 2}}{c_{\mu 2}(c_{\beta 2} - \gamma_1)}$$

These two conditions are also required for the equilibrium with $RS$ at fixation to be stable, in addition to:

$$\frac{\tau}{\mu} < \frac{c_{\beta 1}c_{\mu 1} - c_{\beta 2}c_{\mu 2}}{c_{\mu 1}c_{\mu 2}(\gamma_2 c_{\beta 2} - \gamma_1 c_{\beta 1})}$$

Therefore, the relative treatment rate ($\gamma_1$) and fitness cost parameters at which incomplete linkage disequilibrium may occur are constrained by: $\frac{1-c_{\mu 2}c_{\beta 2}}{c_{\mu 2}(c_{\beta 2}-\gamma_1)} < \frac{c_{\beta 1}c_{\mu 1}-c_{\beta 2}c_{\mu 2}}{c_{\mu 1}c_{\mu 2}(\gamma_2 c_{\beta 2}-\gamma_1 c_{\beta 1})} < \frac{1-c_{\mu 1}c_{\beta 1}}{\gamma_1 c_{\mu 1}c_{\mu 2}c_{\beta 1}}$. In addition, for incomplete linkage equilibrium to actually be observed, there must be at least one stratum with resistance proneness $\frac{\tau}{\mu} > \frac{c_{\beta 1}c_{\mu 1}-c_{\beta 2}c_{\mu 2}}{c_{\mu 1}c_{\mu 2}(\gamma_2 c_{\beta 2}-\gamma_1 c_{\beta 1})}$ and one stratum with resistance proneness $\frac{\tau}{\mu} < \frac{c_{\beta 1}c_{\mu 1}-c_{\beta 2}c_{\mu 2}}{c_{\mu 1}c_{\mu 2}(\gamma_2 c_{\beta 2}-\gamma_1 c_{\beta 1})}$.

This places considerable restrictions on the parameter range at which incomplete linkage equilibrium could be observed. For example, consider a system in which strata correspond to strains differing in clearance rate, with a uniform total antibiotic consumption rate across the strata. For parameter values $c_{\mu 1} = 0.99$, $c_{\mu 2} = 0.9$, $\gamma_1 = 0.09$, $c_{\beta 1} = c_{\beta 2} = 1$ and total antibiotic consumption rate $\tau$ of one prescription per year per person, incomplete linkage equilibrium will be observed only if there is at least one strain with clearance rate $\mu$ between 0.677 and 0.743 per month and another with clearance rate between 0.668 and 0.677 per month - or, equivalently, duration of carriage between 1.35 and 1.48 months and duration of carriage between 1.48 and 1.50 months (mean duration of carriage is $\frac{1}{\mu}$). If these conditions are not met, incomplete linkage disequilibrium, though theoretically possible, will not be observed.

SI Figure 2 and SI Figure 3 further illustrate this sensitivity to parametrisation. In particular, the closer the rate at which the two drugs are consumed, the more restricted the parameter space in which incomplete linkage disequilibrium can arise.

Finally, it is worth noting that when the fitness cost of resistance affects transmission rate only, a similar analysis shows that incomplete linkage disequilibrium never arises. In summary, under specific conditions (the cost of resistance affecting clearance rate and more commonly used antibiotics having a greater fitness cost),

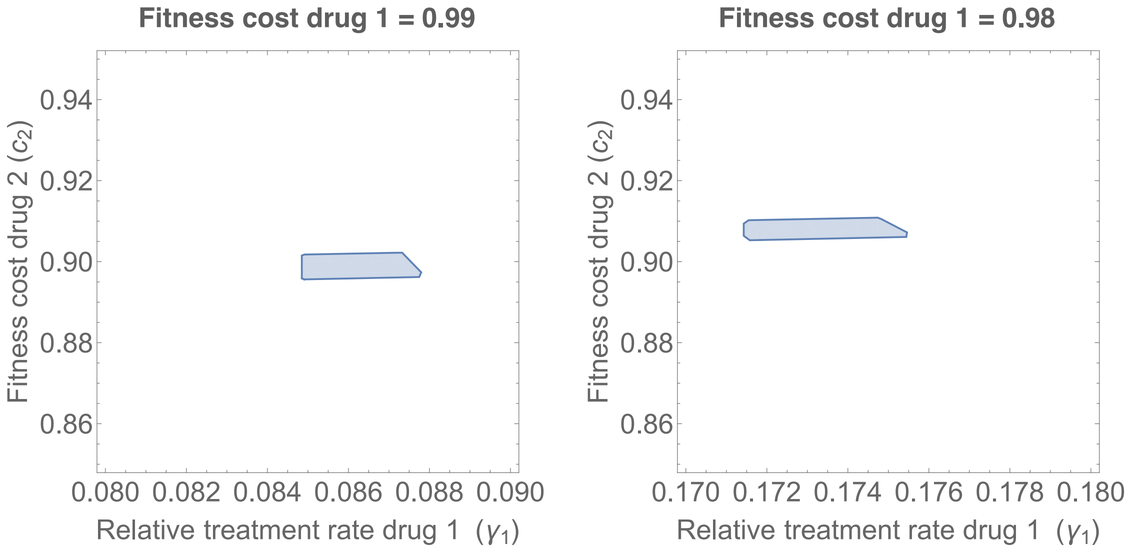


Figure 2: The effect of relative treatment rate ($\gamma_1$) and fitness cost on whether incomplete linkage disequilibrium is observed. The plots assume one stratum with clearance rate $\mu = 0.65$ and another with clearance rate $\mu = 0.70$ and the shaded area represents where $RS$ is selected for in one stratum and $SR$ in the other. Other parameter values are $\tau = 1/12$, $c_{\beta 1} = c_{\beta 2} = 1$ and therefore $c_1 = c_{\mu 1}$ and $c_2 = c_{\mu 2}$. Small changes in the clearance rates of the strata (e.g. to $\mu = 0.60$ and $\mu = 0.65$) lead to complete linkage disequilibrium being observed for all values of $\gamma_1$ and $c_2$. The same is true for increasing the fitness cost of the less commonly prescribed drug (e.g. $c_1 = 0.97$).

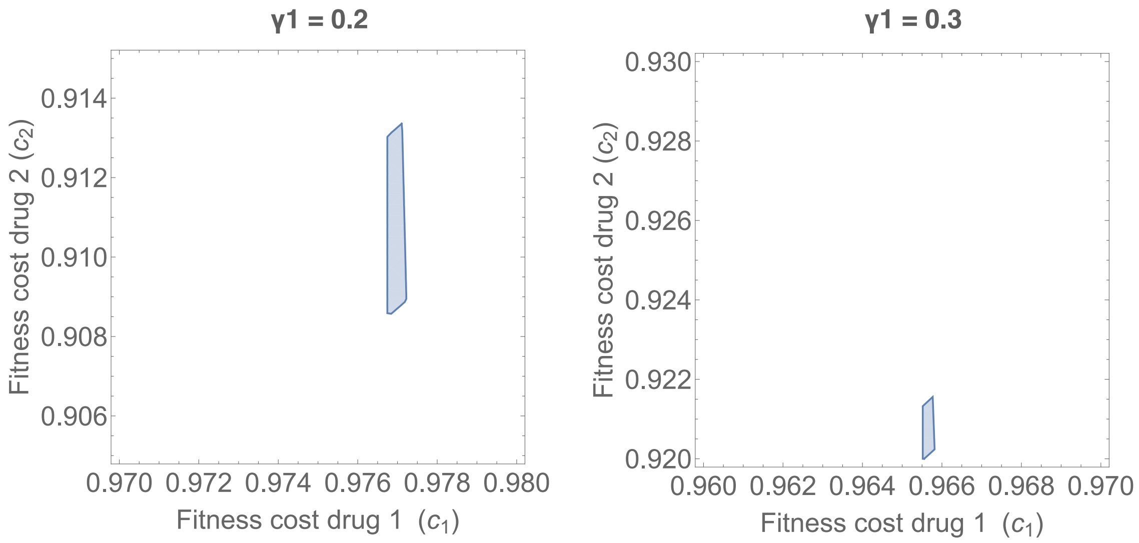


Figure 3: The effect of relative treatment rate ($\gamma_1$) on the range of fitness costs at which incomplete linkage disequilibrium is observed. The plots assume one stratum with clearance rate $\mu = 0.65$ and another with clearance rate $\mu = 0.70$ and the shaded area represents where $RS$ is selected for in one stratum and $SR$ in the other. Other parameter values are $\tau = 1/12$, $c_{\beta 1} = c_{\beta 2} = 1$ and therefore $c_1 = c_{\mu 1}$ and $c_2 = c_{\mu 2}$. At higher values of $\gamma_1$ (e.g. $\gamma_1 = 0.4$), complete linkage disequilbrium in observed for all values of $c_1$ and $c_2$.

incomplete linkage disequilibrium is theoretically possible. However, the narrowness of the parameter range in which this occurs suggests this mechanism leading to incomplete linkage disequilibrium and non-nested resistance profiles being observed is very rare.

### 4 Recombination rate

Allowing recombination at the resistance loci breaks down linkage disequilibrium and therefore decreases the association between resistance determinants: increasing the recombination rate parameter ($r$) in the model represented by Equation 10 in the main text decreases mean linkage disequilibrium (Figure 3 in main text). The $r$ parameter is not directly interpretable as the frequency at which recombinant strains are transmitted: we assume recombination requires co-infection - how often a host infected by any particular strain transmits a recombinant strain therefore also depends on the frequency of other strains. Here, to make values of $r$ more interpretable, we explore its relationship to the proportion of transmission events involving a recombinant strain: $r \sum_x \sum_y I_{xy}$. (Note that this includes all recombination events, even those leading to transmission of an identical genotype).

Mostowy et al. have estimated that the PMEN1 pneumococcal lineage undergoes recombination events (*at any locus*) at a rate of approximately 0.01 events per month [2]. Pneumococcal transmission rates, on the other hand, have been estimated to be around 1.3-4.3 per month [3], suggesting that transmission of recombinant strains, where recombination has occurred at *any* locus, accounts for 0.2 - 0.8% of transmission. The upper bound of the $r$ parameter range explored in the main text ($r = 0.01$), on the other hand, corresponds to transmission of strains where recombination has occurred *specifically at one of the three resistance loci* making up 1.7% of transmission events (SI Figure 4). Some resistance loci in the pneumococcus are recombination hotspots, but even so, this recombination rate is unrealistically high. Values of linkage disequilibrium remain relatively high at this unrealistic rate of recombination (Figure 3 in main text).

It is worth noting, however, that recombination rates for mobile genetic element (plasmid or transposon) associated resistance determinants may be higher if they occur on different elements. Very high rates of recombination would also abolish coexistence - and this effect occurs more rapidly than the break down of linkage disequilibrium (SI Figure 5). Therefore, when fitness variation in the effect of resistance acts as a mechanism of coexistence, it will also produce linkage disequilibrium, even in the presence of recombination.

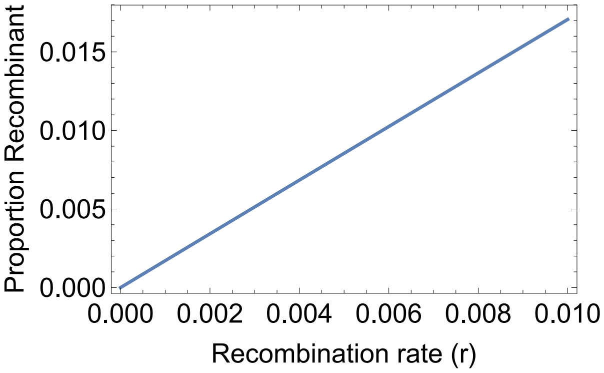


Figure 4: The proportion of transmission of recombinant strains as a function of the recombination rate parameter $(r)$ in the model represented by Equation 10 in the main text.

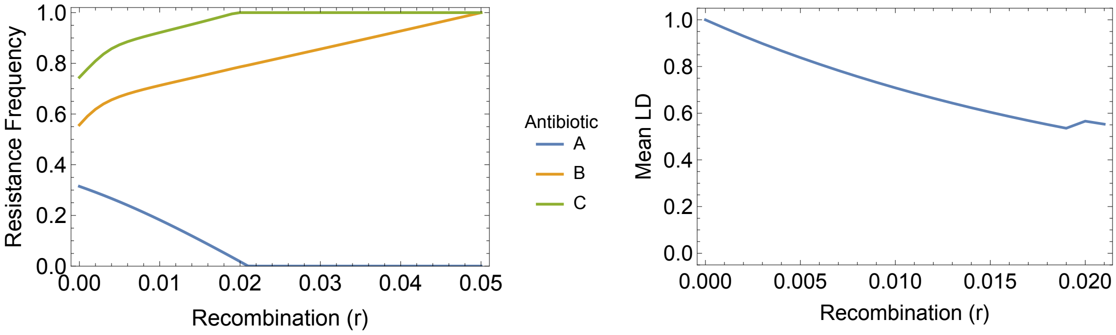


Figure 5: Strain frequencies and mean linkage disequilibrium (LD) between resistances in a model with three antibiotics $(A,\ B$ and $C)$ consumed at different rates and five strains differing in duration of carriage with increasing rates of recombination at the resistance loci (see Methods in main text). Very high rates of recombination eliminate coexistence before the association between resistance determinants. Note the difference in x-axis range: LD is only defined between loci with more than one allele so once coexistence is only observed for a single locus, mean LD can no longer be calculated.
